# TISCH: a comprehensive web resource enabling interactive single-cell transcriptome visualization of tumor microenvironment

**DOI:** 10.1101/2020.08.15.251959

**Authors:** Dongqing Sun, Jin Wang, Ya Han, Xin Dong, Rongbin Zheng, Jun Ge, Xiaoying Shi, Binbin Wang, Ziyi Li, Pengfei Ren, Liangdong Sun, Yilv Yan, Peng Zhang, Fan Zhang, Taiwen Li, Chenfei Wang

**Affiliations:** Clinical Translational Research Center, Shanghai Pulmonary Hospital, School of Life Science and Technology, Tongji University, Shanghai, 200433, China; Department of Thoracic Surgery, Shanghai Pulmonary Hospital, School of Medicine, Tongji University, Shanghai, 200433, China; State Key Laboratory of Oral Diseases, National Clinical Research Center for Oral Diseases, Chinese Academy of Medical Sciences Research Unit of Oral Carcinogenesis and Management, West China Hospital of Stomatology, Sichuan University, Chengdu, Sichuan, 610041, China

## Abstract

Cancer immunotherapy targeting co-inhibitory pathways by checkpoint blockade shows remarkable efficacy in a variety of cancer types. However, only a minority of patients respond to treatment due to the stochastic heterogeneity of tumor microenvironment (TME). Recent advances in single-cell RNA-seq technologies enabled comprehensive characterization of the immune system heterogeneity in tumors, but also posed computational challenges on how to integrate and utilize the massive published datasets to inform immunotherapy. Here, we present Tumor Immune Single Cell Hub (TISCH, http://tisch.comp-genomics.org), a large-scale curated database that integrates single-cell transcriptomic profiles of nearly two million cells from 76 high-quality tumor datasets across 28 cancer types. All the data were uniformly processed with a standardized workflow, including quality control, batch effect removal, malignant cell classification, cell clustering, cell-type annotation, differential expression analysis, and functional enrichment analysis. TISCH provides interactive gene expression visualization across multiple datasets at the single-cell level or cluster level, allowing systematic comparison between different cell-types, patients, tissue origins, treatment and response groups, and even different cancer-types. In summary, TISCH provides a user-friendly interface for systematically visualizing, searching, and downloading gene expression atlas in the TME from multiple cancer types, enabling fast, flexible and comprehensive exploration of the TME.

## INTRODUCTION

Cancer is a leading cause of death worldwide (1). In recent years, cancer immunotherapy has emerged as one of the most promising therapeutic strategies and demonstrated remarkable efficacy in tumor elimination and control (2). One major obstacle for immunotherapy is that only a small fraction of patients can benefit from the treatment due to the highly complex and heterogeneous tumor microenvironment (TME) (3). Therefore, it is of vital importance to investigate the detailed cell-type compositions and characterize gene expression dynamics in TME, which could potentially improve the utility of cancer immunotherapy.

Single-cell RNA sequencing (scRNA-seq) has been increasingly adopted to investigate cell phenotypes, states, functions, and crosstalk in the TME (4). It provides an unprecedented resolution to decipher the heterogeneous populations in TME, allowing identification of novel cell-types and discovery of unknown associations (5). For example, Zheng et al. characterized the infiltrated T-cells of liver cancer using scRNA-seq, and identified *LAYN* as a marker for expanded tumor Treg and exhausted CD8 T-cells (6). Guo et al. discovered a “pre-exhausted” stage of T-cells and bimodal distribution of *TNFRSF9* in Tregs from non-small-cell lung cancer (NSCLC), suggesting previously unknown heterogeneity of the infiltrated T-cells (7). A recent study performed on melanoma patients treated with checkpoint therapy showed that patients with high TCF7^+^CD8^+^ T-cells are associated with positive clinical outcomes after treatment (8). These studies proved that single-cell transcriptomics has enabled the cancer biologists and oncologists to better understand the TME heterogeneity and provided novel clinical implications. However, the rapidly accumulated tumor scRNA-seq data have also posed great computational challenges for data integration and reuse.

There have been efforts to systematically collect and curate single-cell datasets, such as CancerSEA, scRNASeqDB, SCPortalen, PanglaoDB, and JingleBells (9-13). Among them, only CancerSEA is cancer-related, although it only focuses on cancer cells without considering immune or stromal cells in the TME. Moreover, most of these databases contain a limited number of cells. CancerSEA (9) explores the functional heterogeneity of only 41,900 cancer cells, and SCPortalen (11) only has 67,146 cells combining human and mouse datasets. Large scale repositories, such as Single Cell Portal from the Broad Institute (14) and Single Cell Expression Atlas from European Bioinformatics Institute (EMBL-EBL) (15), provide greater numbers of datasets, but they are not cancer-focused, and have limited and often inconsistent cell-type annotations across datasets. So far there are still no comprehensive, intuitive, and convenient web resources with user-friendly interactive features for researchers to explore public tumor scRNA-seq datasets.

Here, we present Tumor Immune Single Cell Hub (TISCH), a comprehensive and curated web resource aiming to decipher the complex components of the TME at single-cell resolution. TISCH builds a scRNA-seq atlas of 76 high-quality tumor datasets across 28 cancer types, which was collected from Gene Expression Omnibus (GEO) and ArrayExpress (see Methods). The TISCH atlas includes nearly two million cells, of which 378K were malignant cells and 1,566K were non-malignant cells. 17 datasets have tumors undergoing immunotherapy treatment (Figure 1), and three additional PBMC datasets from healthy donors were included to provide baseline expression level for immune cells. These datasets were uniformly processed with a standardized workflow, including quality control, batch effect removal, malignant cell classification, clustering, differential expression analysis, curated multi-level cell-type annotation, and functional enrichment analysis. TISCH provides a user-friendly interface to support interactive exploration and visualization of each dataset or across multiple datasets at both single-cell and annotated cluster levels. The continued maintenance and update of TISCH promise to be of great utility to the immuno-oncology community.

**Figure 1.**
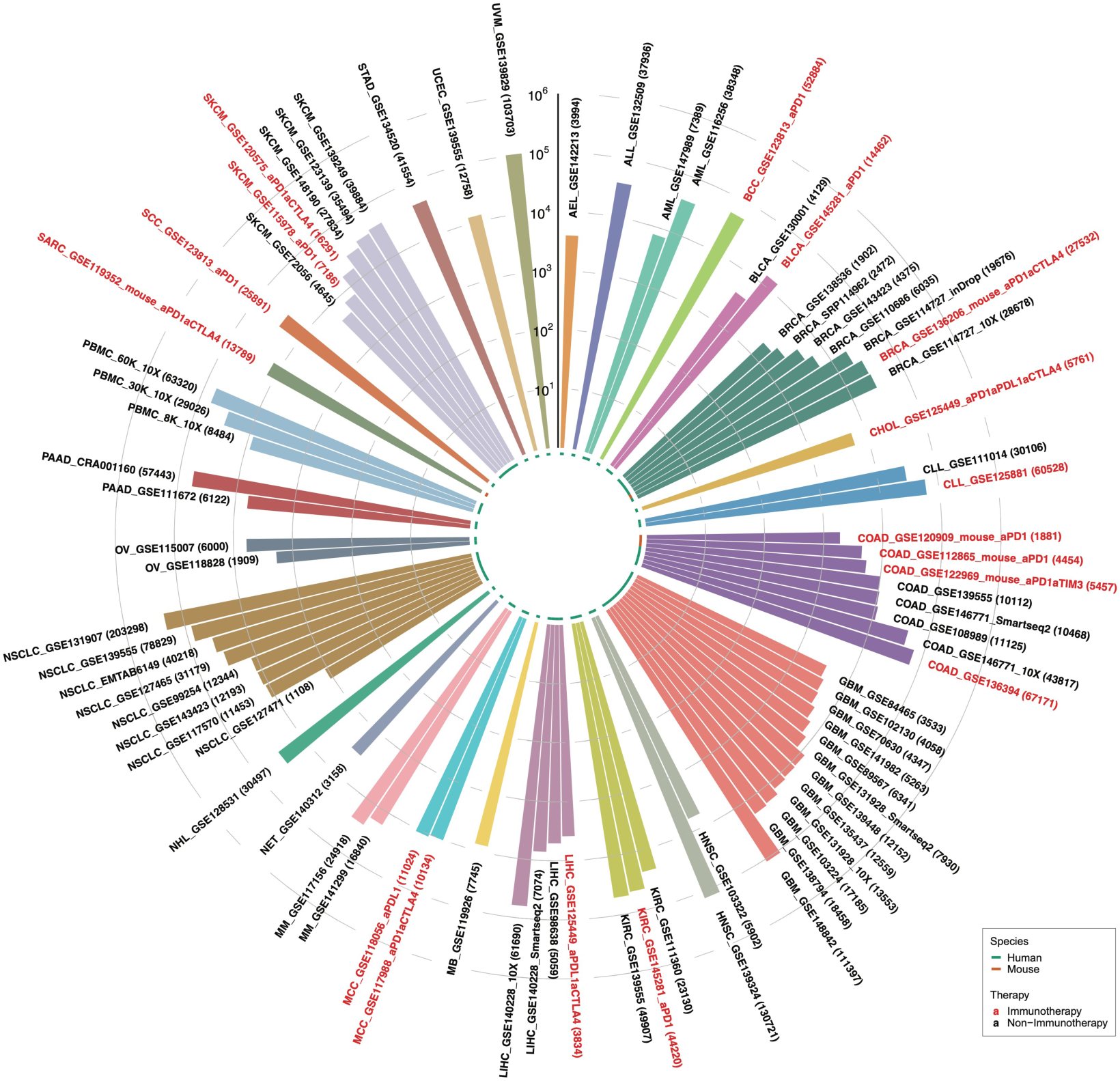
Summary of TISCH datasets. TISCH includes 79 high-quality single-cell datasets, covering nearly 2 million cells across 28 cancer types. Datasets on human and mouse tumors are indicated as green and orange in the inner circle, respectively. Datasets with immunotherapy are labeled in red. The number of cells for each dataset is shown inside the parenthesis.

## MATERIALS AND METHODS

### Data collection and meta information curation

We developed a text-mining-based data parsing workflow and collected tumor scRNA-seq datasets of human from GEO and ArrayExpress. We searched the single-cell keywords such as ‘single cell RNA sequencing’ or ‘scRNAseq’ or ‘single cell’ or ‘single-cell’, as well as the technology-related keywords like ‘microfluidics’, ‘10X Genomics’ and ‘SMARTseq’, and the tumor-related keywords such as ‘tumor’ or ‘cancer’ or ‘carcinoma’ in the description page of GEO or ArrayExpress. Each dataset was then manually confirmed and curated. A total of 118 cancer-related scRNA-seq datasets were obtained initially. We kept the datasets with more than 1000 high-quality cells. To expand the utility of TISCH, we also included the scRNA-seq datasets of mouse treated with immunotherapy and three scRNA-seq datasets of human peripheral blood mononuclear cells (PBMC) from 10X Genomics. Overall, the TISCH database contains 76 high-quality tumor datasets across 28 cancer types and three PBMC datasets (Supplementary Table S1). For each dataset, we downloaded the expression matrix of the raw count, TPM and FPKM (if available), and collected sample information from databases or the original paper, such as the patient ID, tissue origin, treatment condition, response groups, and the original cell-type annotation.

### Data Pre-processing

We applied a standardized analysis workflow based on MAESTRO v1.1.0 (16,17) for processing all the collected datasets, including quality control, batch effect removal, malignant cell classification, cell clustering, differential expression analysis, cell-type annotation and gene set enrichment analysis (GSEA) (Figure 2). The quality of cells was determined by two metrics: the number of total counts (UMI) per cell (library size) and the number of detected genes per cell. Low-quality cells were filtered out if the library size was less than 1000 or the number of detected genes was less than 500 (Supplementary Figure S1A). We evaluated the potential batch effect between different patients, treatment conditions, and tissue origins, and removed the batch effect by canonical correlation analysis (CCA) (18) (Supplementary Figure S1B). If a dataset contained multiple cancer types, each cancer type was processed individually. The source code for processing all the collected scRNA-seq dataset are deposited at the Github repository (https://github.com/DongqingSun96/TISCH/tree/master/code).

**Figure 2.**
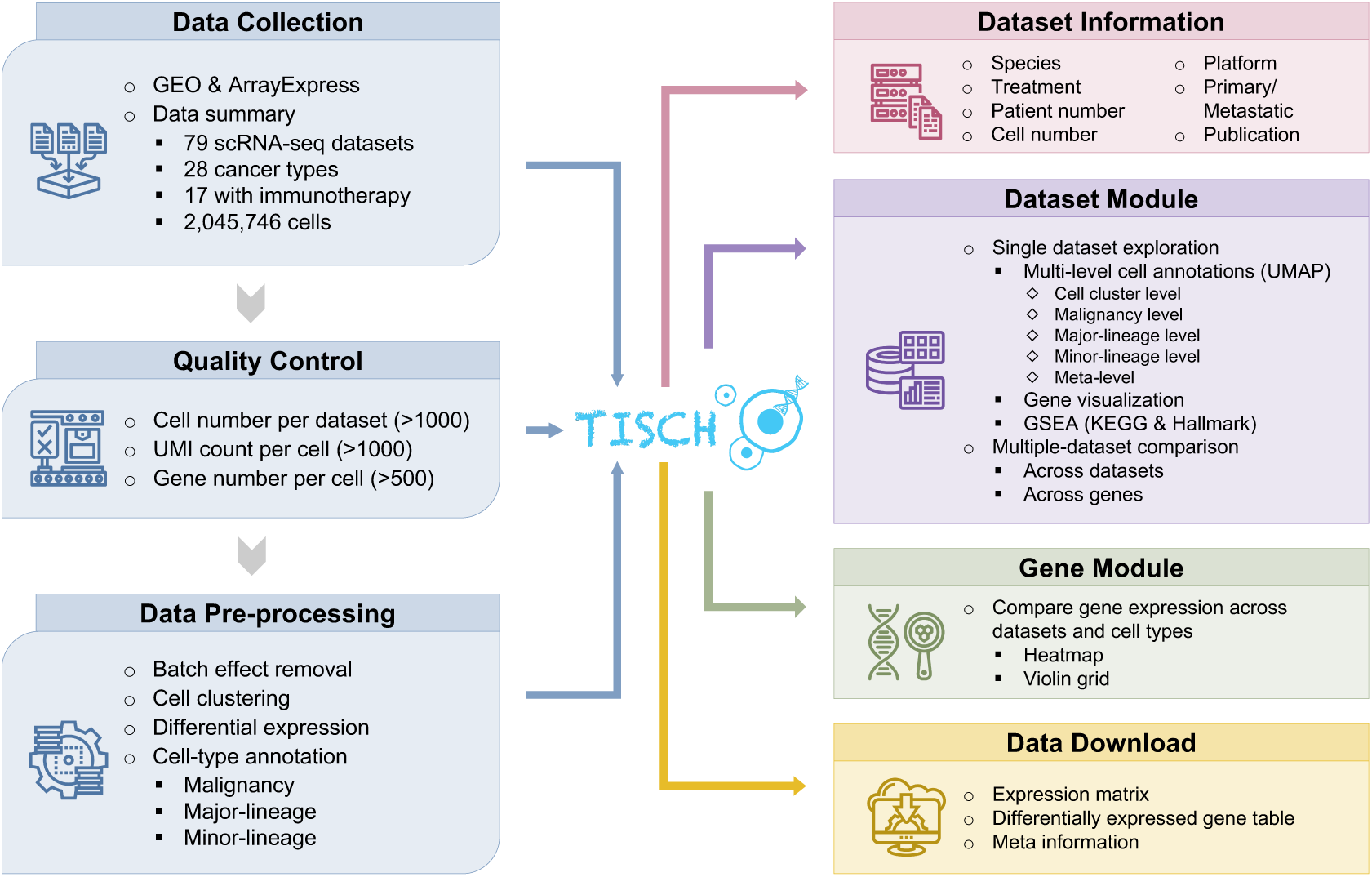
Overview of the TISCH workflow and features. TISCH automatically parsed and curated tumor single-cell RNA-seq datasets from GEO or Array Express databases. All datasets were then uniformly processed with a standardized workflow, including quality control, batch effect removal, cell clustering, differential expression analysis, cell type annotation at multiple levels. Each dataset in TISCH is displayed with relevant study information, including species, treatment, the number of patients and cells, technology platform, stage, and related study. In the Dataset module, TISCH provides two functions: single-dataset exploration and multiple-dataset comparison. In the Gene module, TISCH allows single gene expression visualization across multiple datasets and cell types. TISCH also supports downloading of expression matrices, DE gene tables and meta-information for each dataset.

### Cell Clustering and Differential Gene Analysis

For each dataset, the MAESTRO workflow identified the top 2000 variable features, and employed PCA for dimension reduction, KNN, and Louvain algorithm to perform cell clustering (19,20). To better capture the cellular difference and variabilities for datasets with different cell numbers, we adjusted the number of principal components and the resolution for graph-based clustering, which were both increased with the cell number (Supplementary Table S2). The uniform manifold approximation and projection (UMAP) were utilized to further reduce the dimension and to visualize the clustering results (21). We applied the Wilcoxon test to identify differentially expressed (DE) genes of each cluster compared to all other cells based on the log-transformed fold change (|logFC| >= 0.25) and false discovery rate (FDR < 1e-05).

### Cell-type annotation

The clusters of malignant cells were determined by combing three approaches. First, we took the cell-type annotations provided by the original studies. Second, we ran InferCNV v1.2.1 (22) to predict cell malignancy based on the predicted copy number variation and separated the cells into malignant and non-malignant clusters. Third, if available, we scored the expression level based on the malignant marker genes from the original studies (Supplementary Figure S1C). For the other normal clusters, we automatically annotated the cell clusters based on the DE genes by improving the marker-based annotation method in MAESTRO. The marker genes of each cell type were collected from the published articles (23-25) and curated manually (Supplementary Table S3, S4). In each cluster, we calculated the average logFC of the marker genes for each cell type and took it as cell-type score *S*_*c*_. Finally, each cluster will be assigned a specific cell type *C*_*j*_, which has the highest score among all cell types.

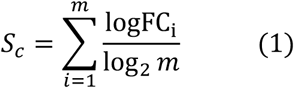

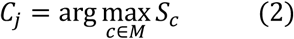

Where *M* is the set of all collected cell types, *m* is the number of marker genes for a certain cell type *c* in *M*. logFC_i_ is the logFC of marker gene *i* in cell type *c*.

To optimize the capacity of the marker-based cell-type annotation, we created a parameter cutoff for 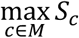 and set the default value to 0.06 based on nine datasets with original cell-type annotation. The automatic cell-type annotation 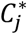 is predicted as:

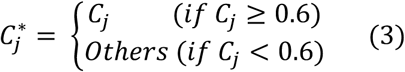

Consequently, we retained 18 common cell types at the major-lineage level, such as B cells, CD8^+^ T cells (CD8T), CD4^+^ T cells (CD4Tconv) (Supplementary Figure S2 and Supplementary Table S3). To gain more detailed insights into immune cells, we collected and curated the cell subtype signatures from the public literature (Supplementary Figure S2 and Supplementary Table S4), and we further generated the annotation of minor-lineage subtypes differentiating from major-lineage cell-types. For example, typical CD8+ T cells at the major-lineage level could differentiate into naïve CD8^+^ T cells (CD8Tn), central memory CD8^+^ T cells (CD8Tcm), effector memory CD8^+^ T cells (CD8Tem) and effector CD8^+^ T cells (CD8Teff). After automatic cell-type annotation, we made manual corrections to some tissue-specific cell types by combining them with original annotation and malignant cell identification in the previous step. All the cells were classified into three types, malignant cells, immune cells, and stromal cells based on the major-lineage level annotation, and the annotations were defined as the malignancy level (Supplementary Figure S2).

### Functional enrichment analysis

To characterize the functions of distinct cell-type populations, we performed gene set enrichment analysis (26,27) according to the rank of genes based on the fold-change from the differential analysis. We collected 186 gene sets of Kyoto Encyclopedia of Genes and Genomes (KEGG) pathways (28) and 50 of hallmark pathways from the Molecular Signatures Database (MSigDB v7.1) (29). Significant up-regulated and down-regulated pathways (FDR <= 0.05) in each cluster were identified and visualized to enable the function comparison between clusters in each cell type. In addition, for the datasets with treatment, functional enrichment analysis of each cell type between different treatment conditions was also performed if the treatment information was available. This analysis was fulfilled by GSEA v4.0.3 for Linux, and figures were generated by the ComplexHeatmap R package v1.99.5 (30).

### Gene conversion

To enable the consistent gene format across different assemblies and species, genes of each human and mouse dataset were converted into GRCh38.p13 and GRCm38.p6, respectively. Then, the homologous relations between GRCh38 and GRCm38 genes were constructed through ‘getLDS’ function of biomaRt package v2.42.0 (31) to support gene search across species in TISCH. For those genes with one-to-many relations between species, only one homology mapping was retained randomly.

### Gene visualization across cancer types and cell types

Datasets with a large number of cells (>10000) will usually consume high memory and take long response time to generate expression visualization figures. To ensure the quick response for users when searching a gene across multiple cancer types and cell types, we applied a sub-sampling procedure for 49 datasets with more than 10,000 cells. For each gene, we sorted the cells according to the expression level of the gene in each cluster with more than 200 cells. Every 10 cells were assigned into a bin and the median of the 10 cells was calculated to represent the expression level of the bin. For clusters with less than 200 cells, all the cells were kept directly. Each point in the gene expression violin plot represents a bin and the distribution of bins was showed between different cell-types and datasets. This method collapsed large datasets into almost one-tenth of the original ones, and significantly improve the speed of read-in and generating the gene expression visualization figures.

### Web portal for the database

Based on the uniformly processed scRNA-seq datasets, we build the TISCH web portal to present the analysis results in a user-friendly way. All the processed and annotated datasets can be searched, visualized, and downloaded from the web portal. The front-end display is achieved through HTML and CSS, and the back-end data are organized and queried by MySQL database management system v8.0.20. The interaction between the front-end and back-end is enabled through JavaScript and Python. All the charts in TISCH are generated by Highcharts v8.1.2 and in-house Python and R scripts. TISCH database is deployed with Apache2 HTTP server and is freely available at http://tisch.comp-genomics.org without any registration or login. All the functions of TISCH have been tested in Google Chrome and Apple Safari browsers.

## RESULTS

### Dataset summary in TISCH

The current TISCH database contains a total of 2,045,746 cells from 79 datasets involving 28 cancer types, with 378,392 malignant cells and 1,667,354 non-malignant cells. In TISCH, there are 76 tumor-related datasets, including 17 tumor datasets with immunotherapy treatment (12 human datasets and 5 mouse datasets) (Figure 1). Three additional PBMC datasets from healthy donors are included to provide baseline expression levels for immune cells. On average, each dataset has 26,455 cells, with one largest dataset from NSCLC have over 200K cells (Supplementary Table S1). In total, TISCH covered 68,287 genes for human datasets and 18,789 genes for mouse datasets, with an average of 18,411 genes covered per dataset.

### Utility of TISCH

TISCH presents all the analysis results including clustering, differential gene identification, cell-type annotation, and GSEA in a user-friendly interface for public accessing. TISCH provides two modules for users to visualize the datasets (Figure 2). The Dataset module supports the detailed exploration of an individual dataset. In addition, it also supports multiple gene expression visualizations across multiple datasets at the single-cell level. The Gene module allows single gene visualization across multiple different scRNA-seq datasets at the cell-type level.

### Single-dataset exploration

In the Dataset module, TISCH supports the advanced search for datasets of interest to explore the cell-type composition, gene expression distribution, functional status of each cell-type, and comparison between different tissue-origin or treatment groups. If users focus on one specific cancer type, they can click the corresponding tissue icon on the Home page to query related datasets. In the forwarding Dataset page, users can further narrow down the query results according to other criteria, such as species, treatment, and included cell-types. The datasets satisfying the criteria will be displayed with relevant study information, including the number of patients and cells, technology platform, treatment, stage, and related study.

For each scRNA-seq dataset, the pre-analyzed results of the dataset will be shown in four different tabs, including the overview, gene, GSEA, and download tabs. In the overview tab (Figure 3A), two UMAP plots with cells colored by the cell clusters and cell-type annotations will be displayed on the top. TISCH allows users to choose cell-type annotations from three levels, malignancy level, major-lineage level, and minor-lineage level (Supplementary Figure S2, see Methods). In addition, other meta information, such as patient information, tissue origin, treatment condition and cell-type annotation from the original study can also be displayed if available. On the bottom of the overview page, the top differentially expressed genes for each cluster are provided for users to discover the potential markers of each cell-type. We also allow users to search interested genes and see their relative logFC in different cell-types. In the gene tab (Figure 3B), TISCH provides a gene visualization function to search and compare multiple genes of interest simultaneously in the current dataset. UMAP plots that reflect the expression level of input genes at the single-cell resolution will be returned, enabling the exploration of the co-expression or mutually exclusive relationship between different genes. Besides, a violin plot will be displayed to show the distribution of the interested gene expression in different cell types. TISCH allows users to compare the expression of genes between different groups, such as tissue origins, treatment conditions, or response groups if the meta-information is available (Figure 3B, Supplementary Figure S3A). The statistical significance between different groups was evaluated using Mann-Whitney test for two groups or Kruskal-Wallis test for three or more groups (Figure 3B). In addition to individual gene input, TISCH supports gene list upload so that users can explore the expression pattern of their interested gene signatures at both single-cell and cell-type level. Genes in the uploaded signature list will be collapsed by the mean or median of expression, which depends on users’ choices. In the GSEA tab (Figure 3C), the pre-calculated GSEA results are available for users to characterize the functional differences between different cell-types. Heatmaps will be shown to display the enriched up- or down-regulated KEGG and hallmark pathways identified based on differential genes in each cluster. For the datasets with treatment information, TISCH also provides GSEA results for comparing functional pathways between different treatment conditions or treatment responses for each cell-type.

**Figure 3.**
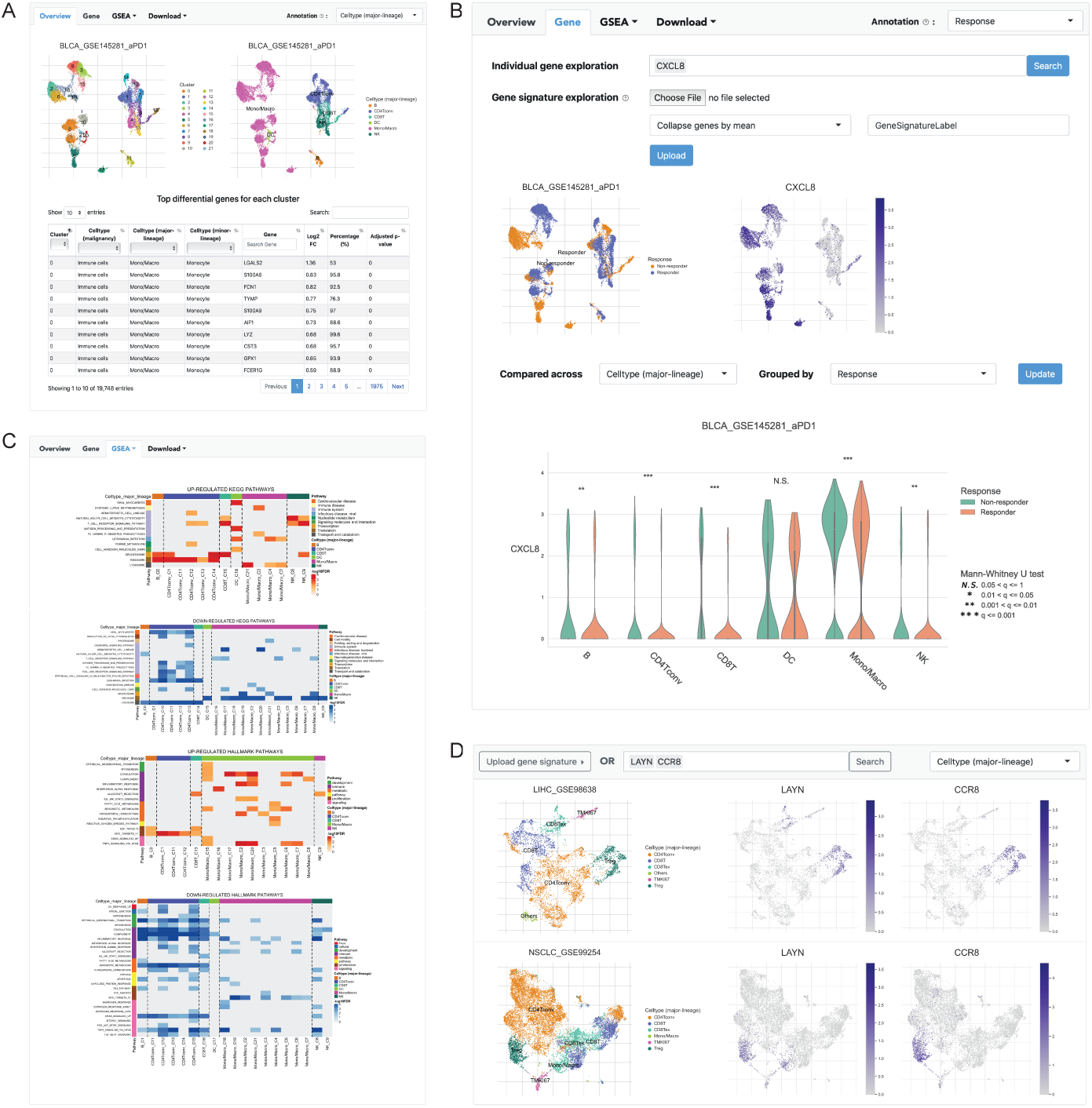
Dataset module of TISCH. (**A**) The overview tab of the BLCA_GSE145281_aPD1 dataset. Two UMAP plots with cells colored by cluster ID (left) and cell type (right) are displayed on the top of the tab. The table below shows DE genes in each cluster. (**B**) The gene tab of the single-dataset module where expression of genes of interest can be visualized at single-cell and cell-type resolution. Two UMAP plots are to show the cell distribution of treatment response groups (left) and the expression of *CXCL8* (right). Comparison of *CXCL8* expression between “Responder” (orange) and “Non-responder” (green) across cell types is visualized by the violin plot. The significance of difference between two groups in each cell type is evaluated through Mann-Whitney U test and adjusted through Banjamini-Hochberg correction. ‘N.S.’ represents q (adjusted p-value) > 0.05, ‘*’ represents 0.01 < q <= 0.05, ‘**’ represents 0.001 < q <= 0.01, and ‘***’ represents q <= 0.001. (**C**) GSEA results of a single dataset. The enriched up- or down-regulated KEGG and hallmark pathways in each cluster are visualized in heatmaps. (**D**) Multiple-dataset module, in which users can compare the gene expression across datasets at single-cell resolution. An example is presented to display the expression of *LAYN* and *CCR8* at single-cell resolution in LIHC_GSE98638 and NSCLC_GSE99254.

Besides the online search and visualization for each dataset, TISCH provides an easy way to download the data containing expression profiles, DE genes, and related meta information. The cluster- or cell-type-averaged expression matrices are archived in a compressed file and can be downloaded by users. The top differential genes of each cluster displayed in the overview tab can also be downloaded. What’s more, TISCH provides three levels of cell-type annotations as well as curated meta-information at the single-cell resolution for downloading. All the figures shown on the web page can also be downloaded in high resolution. Users can utilize the downloaded data for further customized exploration.

To demonstrate an example of exploring the single-dataset module, we queried by caner type “BLCA (Bladder Urothelial Carcinoma)” and focused on the BLCA_GSE145281_aPD1 dataset with anti-PD1 treatment for further analysis. Studies have shown that the difference in patient’s TME may lead to a distinct immunotherapeutic outcome (8,32), we thus compared the different abundance of the cell-type population between responder and non-responder groups. We observed that a higher proportion of monocytes or macrophages are presented in the TME, with apparently more monocytes or macrophages in non-responders (Figure 3A, B). A previous study indicates that *CXCL8*, a major mediator of the inflammatory response, is highly expressed in myeloid cells than lymphoid cells, as well as in non-responders than responders (32). We confirmed this conclusion on BLCA_GSE145281_aPD1 dataset (Figure 3B). Interestingly, similar high expression of *CXCL8* in non-responders’ monocytes or macrophages was also observed in an independent melanoma cohort SKCM_GSE120575_aPD1aCTLA4 (8) (Supplementary Figure S3A). Further GSEA showed that the high expression of *CXCL8* in myeloid cells is associated with down-regulation of the antigen-presentation pathway in both independent datasets, suggesting *CXCL8*-mediated myeloid inflammation might suppress the anti-tumor immunity (32) (Supplementary Figure S3B, C). Hence, this single-dataset module enables quick and interactive gene expression visualization between different cell-types and treatment conditions.

### Multiple-dataset comparison

In addition to single-dataset visualization, TISCH can also facilitate a comparative analysis of multiple datasets at single-cell resolution to explore the potential expression heterogeneity or homogeneity across multiple cohorts. Users can select multiple genes from multiple datasets, and compare the cell-type distribution and gene expression patterns simultaneously (Figure 3D). Similar to single-dataset exploration, TISCH also allows the uploading of gene lists to visualize the averaged expression distribution of candidate gene signatures.

Here we use an example to demonstrate the usage of the multiple-dataset module. It has been reported that *LAYN* and *CCR8* are highly expressed in tumor-infiltrating Treg cells from colon cancer, non-small cell lung cancer, and liver cancer (6,33). We observed the consistently high expression of *LAYN* and *CCR8* in Treg cells from four independent datasets (LIHC_GSE98638, NSCLC_GSE99254, COAD_GSE108989, and COAD_GSE146771_Smartseq2) (6,7,23,34), suggesting the tumor homogeneity in terms of cell phenotype signatures (Figure 3D, Supplementary Figure S4). Besides the Treg cells, *LAYN* is also expressed in a subset of exhausted CD8T cells (Figure 3D, Supplementary Figure S4). As *LAYN* has been linked to immune suppressive function of tumor-infiltrating Treg and exhausted CD8T cells, this indicates the exhausted CD8T cells in the TME are highly heterogeneous and maybe in different exhaustion stage (6). Collectively, the comparative analysis of user-defined features across multiple datasets at single-cell resolution will provide a more detailed and comprehensive insight into the cell-type compositions and gene expression relationships in the TME.

### Gene search across datasets

Although the Dataset module provides a detailed expression distribution for single or multiple datasets, it is often required to quickly locate which cell-type expresses the gene of interest across multiple tumor cohorts and different cancer types. In the Gene module, TISCH provides two ways of visualizing the gene expression from multiple cohorts (Figure 4A). The heatmap displays the input gene expression at the cell-type averaged level (Figure 4B), while the grid violin plot reflects the expression distribution of the input gene at single-cell or 10-cell-binned resolution (Figure 4C). The row names (i.e., dataset names) of the heatmap and the violin plot can be clicked to link to the corresponding single-dataset page, where users can browse and search to achieve a deep understanding of the dataset as described in the single-dataset module.

**Figure 4.**
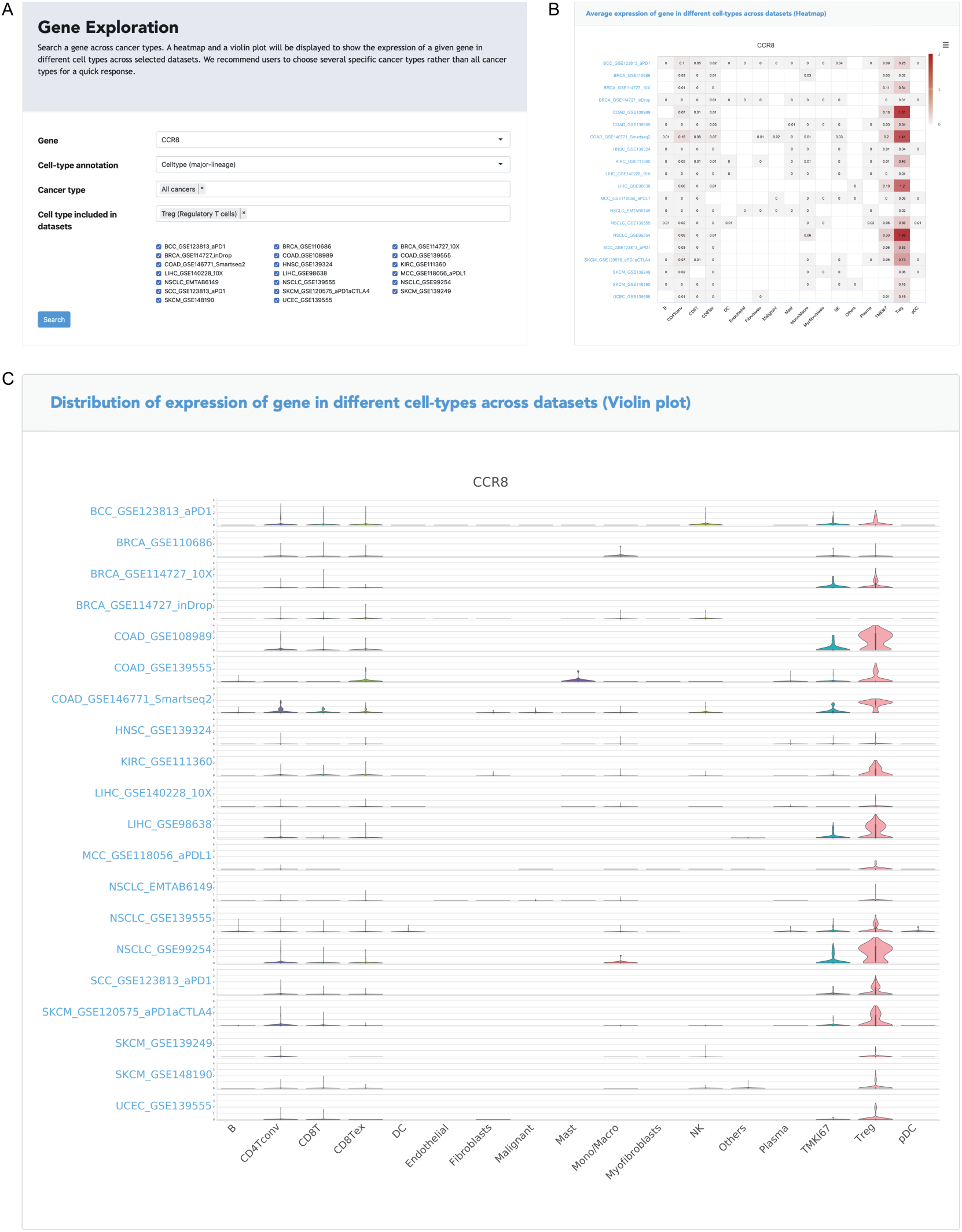
Gene module of TISCH. (**A**) *CCR8* gene search across all cancer types and species. (**B**) The heatmap is to show the expression of *CCR8* in different cell types across all datasets with Treg cells. The color indicates the expression level of the gene. (**C**) The grid violin plot reflects the distribution of gene expression in different cell types across all datasets with Treg cells.

In the previous multiple-dataset module, we have already shown that *CCR8* exhibits cell-type-specific expression in Treg cells from the colon, non-small cell lung, and liver cancer TMEs. It is not clear whether *CCR8* is expressed in other cell types or other cancer types. From the Gene module analysis, it is explicitly observed that *CCR8* also shows highly specific expression in Treg cells for multiple other cancer types, such as melanoma, kidney, and squamous cell carcinoma (Figure 4B). In addition, we observed a bimodal distribution of *CCR8* expression in tumor-infiltrating Tregs cells from multiple cohorts, which is either due to the high drop-out rate of the scRNA-seq dataset, or caused by the heterogeneity within the Treg cells (Figure 4C). Therefore, the Gene module not only empowers the quick location of a specific gene expression pattern across different cell-types, but also helps researchers build a holistic picture of gene expression atlas among different cohorts and cancer-types.

## DISCUSSION

Cancer immunotherapy has brought a paradigm shift to cancer treatment in recent years. Although numerous scRNA-seq datasets have been generated to decipher the complex cell-type compositions and expression heterogeneity in the TME, a well-curated and uniformly processed and annotated data portal for TME scRNA-seq data re-use is still not available. In this context, we present TISCH as a comprehensive single-cell web portal for cancer biologists to investigate and visualize single-cell gene expression in the TME. TISCH shows several advantages compared to the existing single-cell tumor resources. First, TISCH is the most comprehensive TME single-cell data portal to our knowledge, which includes single-cell transcriptome atlas of around 2 million cells from 28 cancer types. The diverse cell types and cancer types present in TISCH enable the investigation of TME heterogeneity in a systematic and holistic view. Second, all the datasets in TISCH were uniformly processed, annotated, and manually curated, which removes the barriers for cross-study comparisons and benefits the data-reuse. Finally, with the meta-information provided, TISCH allows comparisons between different patients, immunotherapy treatment groups, and response groups, showing potential clinical indications for cancer therapy.

In summary, TISCH is a useful repository for TME single-cell transcriptomic data and provides a user-friendly web resource for interactive gene expression visualization of cellular differences across multiple datasets at the single-cell resolution. TISCH will be a valuable resource for cancer biologists and immuno-oncologists to study gene regulation and immune signaling in the TME, identify novel drug targets, and provide insights on therapy response. In the future, we will continue to pay efforts to improve TISCH. We will maintain the web resources regularly to integrate new datasets. We will also provide novel functions in TISCH such as inferring gene-gene co-expression and cell-cell interactions based on expression correlations at the single-cell level. As the increasing numbers of public TME scRNA-seq data are available, we anticipate continued development and maintenance of the TISCH web resource will benefit the broader cancer research community.

## Acknowledgements

This work was supported by the National Natural Science Foundation of China (31801059, 81972551, 81702701). The authors acknowledge X. Shirley Liu and Zexian Zeng from Dana Farber Cancer Institute for the helpful discussion and suggestions on the TISCH website. The authors acknowledge the authors from published studies to share their data on tumor profiling cohorts.

