## Supplementary File for "TISCH: a comprehensive web resource enabling interactive single-cell transcriptome visualization of tumor microenvironment"

### Supplementary Figures
### Figure S1

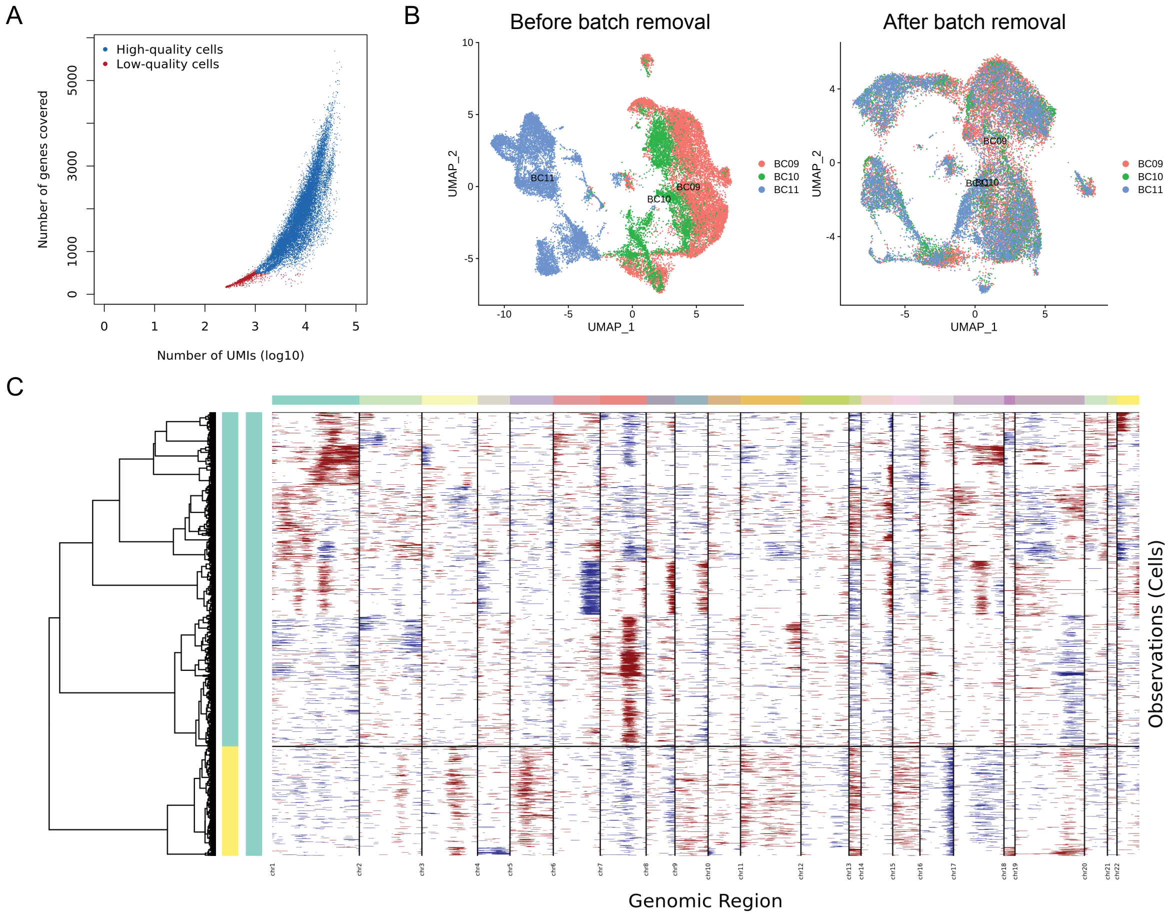


**Data pre-processing.** (A) Quality control for the dataset of MM_GSE141299. The cells with more than 1000 UMI count and 500 gene count are assigned as "High-quality cells" (blue), otherwise "Low-quality cells" (red). (B) Batch effect removal for the dataset of BRCA_GSE114727_10X. The cells across patients are shown before batch removal (left) and after batch removal (right). (C) InferCNV for malignant cell classification for the dataset of MM_GSE141299. Copy number variations are indicated as gain (red) and loss (blue). The left bar represents the malignant cell (green) and non-malignant cell (yellow).

**Figure S2**

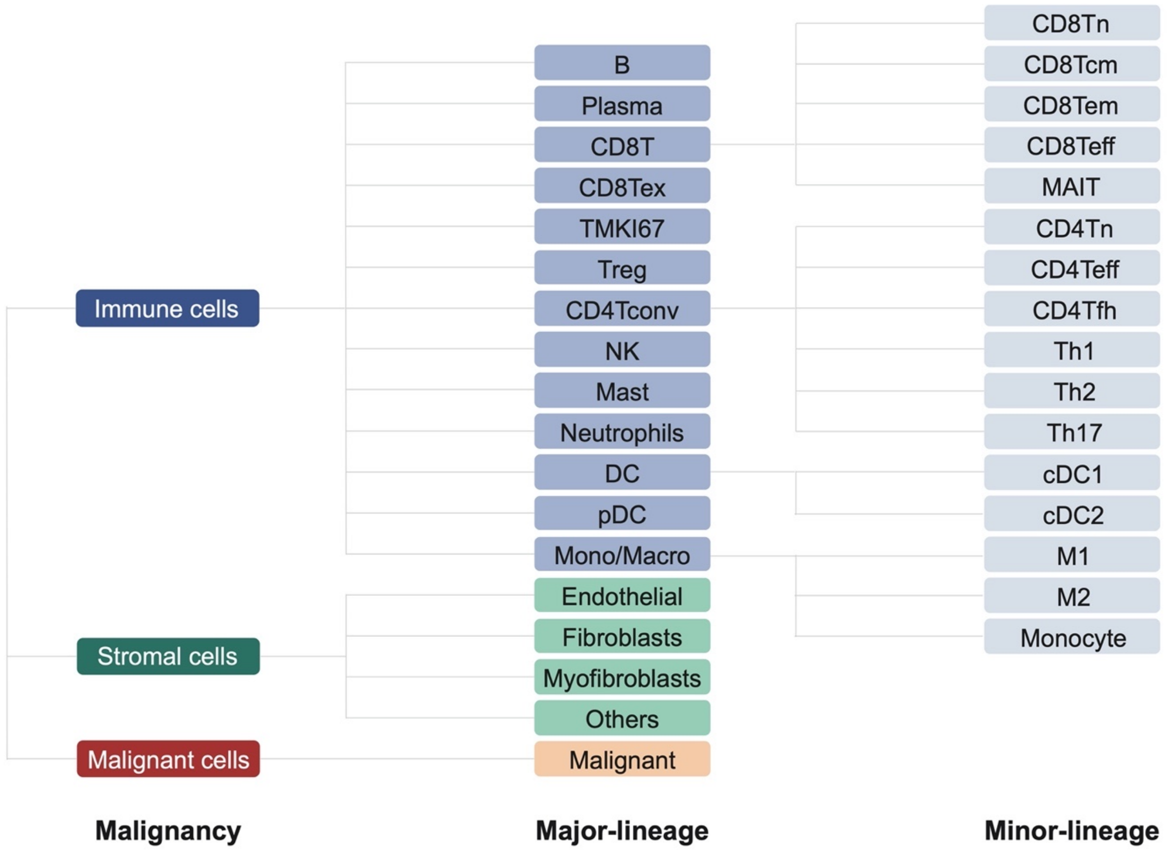


**The hierarchical structure of cell-type annotation.** Cell types are annotated at three levels, including malignancy level (malignant cells, stromal cells, and immune cells), major-lineage level, and minor-lineage level.

### Figure S3

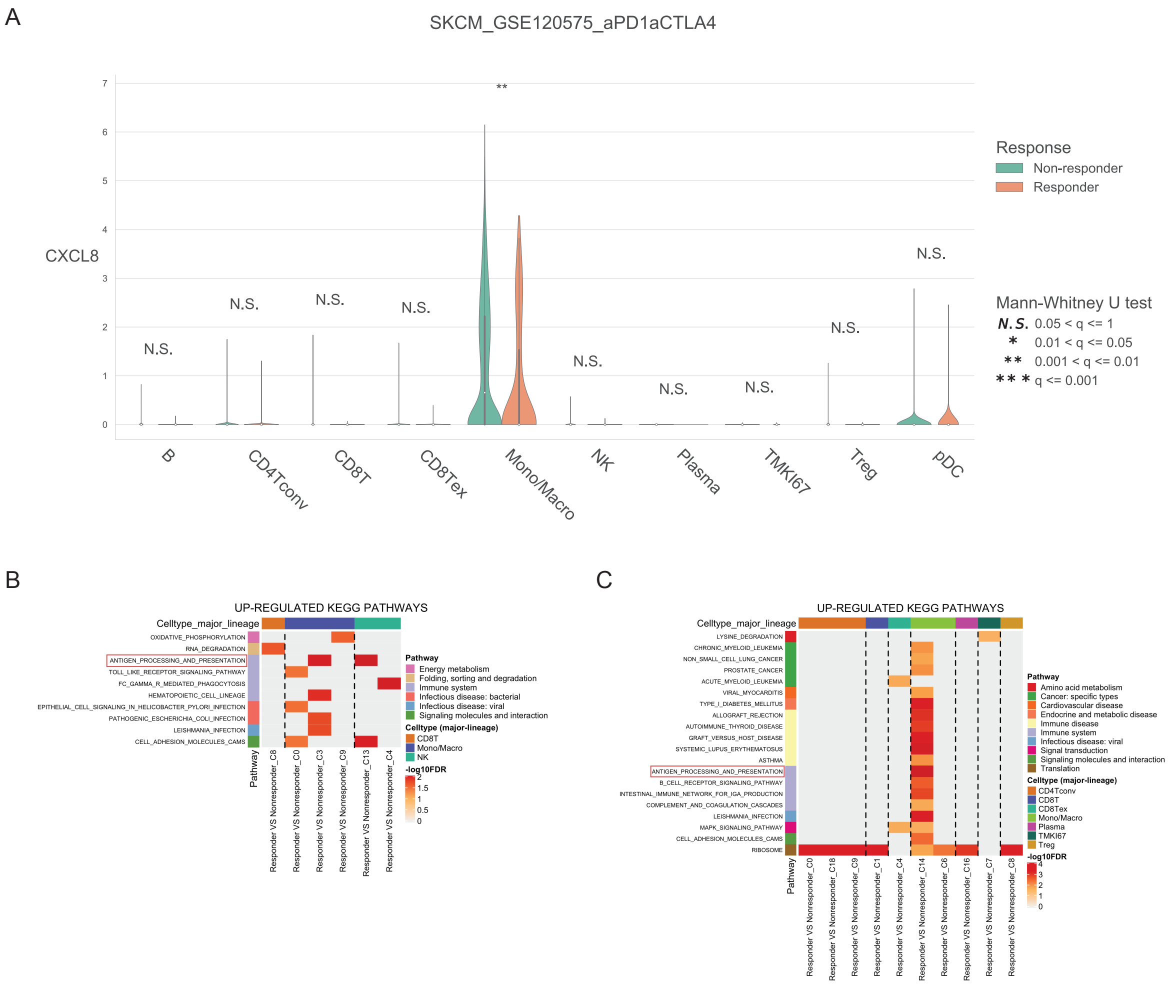


**Example of Dataset module.** (A) The violin plot is to compare the expression of *CXCL8* between 'Responder' (orange) and 'Non-responder' (green) across different cell types in an individual cohort SKCM_GSE120575_aPD1aCTLA4. The significance of difference between two groups for each gene in each cell type is evaluated through Mann-Whitney U test and adjusted through Banjamini-Hochberg correction. 'N.S.' represents q (adjusted p-value) > 0.05, '*' represents 0.01 < q <= 0.05, '**' represents 0.001 < q <= 0.01, and '***' represents q <= 0.001. (B) The enriched up-regulated KEGG pathways for responders in two dependent cohorts BLCA_GSE145281_aPD1 and (C) SKCM_GSE120575_aPD1aCTLA4. ANTIGEN_PROCESSING_AND_PRESENTATION pathway is highlighted in red box.

### Figure S4

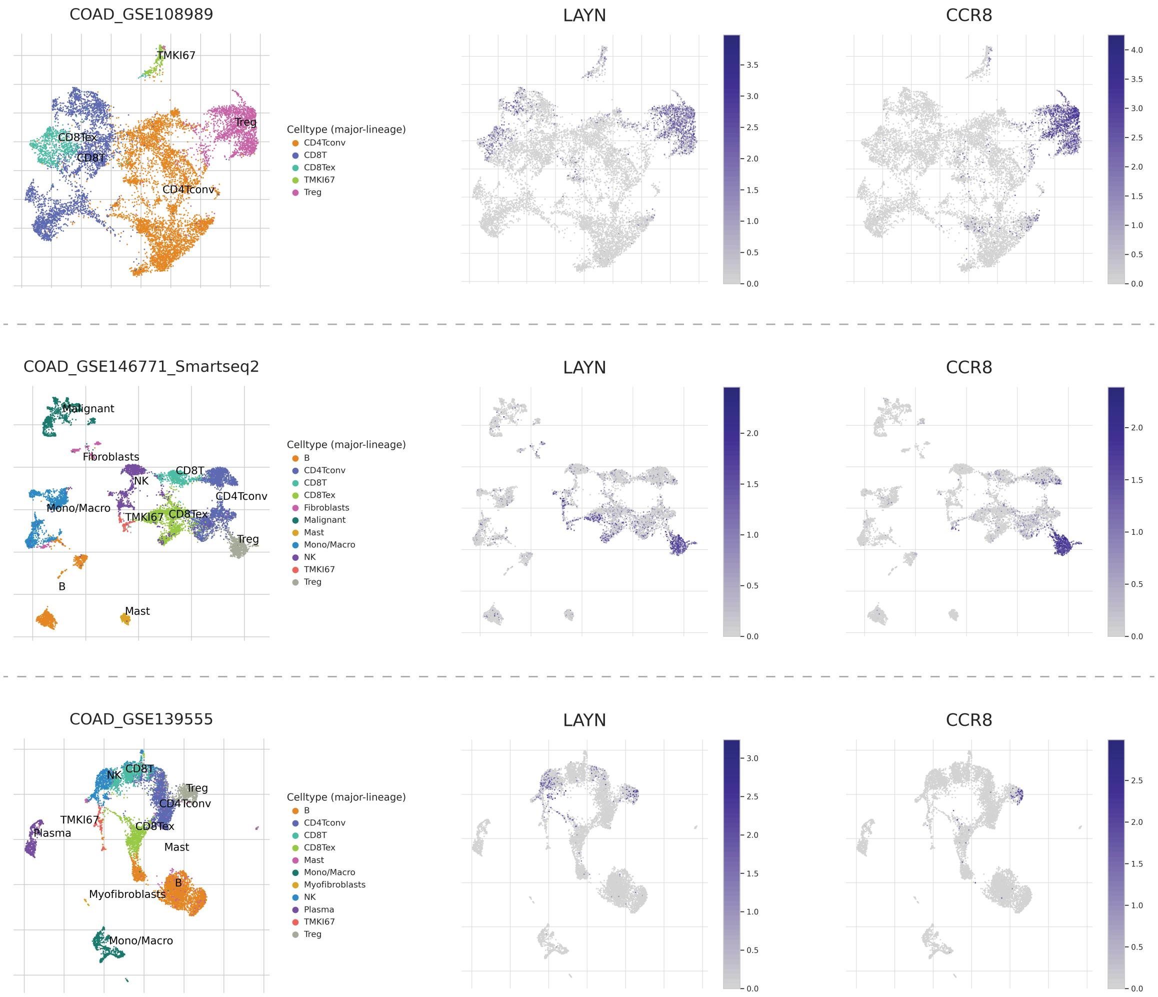


**Comparison of *LYAN* and *CCR8* expression at single-cell resolution across multiple colon cancer cohorts** (COAD_GSE108989, COAD_GSE146771_Smartseq2, and COAD_GSE139555).

**Supplementary Table S1**

The collected data information. TISCH includes 79 high-quality scRNA-seq datasets. The detailed information for each dataset is listed in the table, including dataset name, species, treatment, patient number, cell number, platform, primary/metastatic status, and publication.

| Dataset Name | Species | Treatment | Patients | Cells | Platform | Primary/metastatic | PMID |
|---|---|---|---|---|---|---|---|
| AEL_GSE142213 | Human | TME | 2 | 3994 | 10x Genomics | Primary | 32330454 |
| ALL_GSE132509 | Human | TME | 11 | 37936 | 10x Genomics | Primary | 32415257 |
| AML_GSE116256 | Human | TME | 21 | 38348 | Smart-seq2 | Primary | 29740158 |
| AML_GSE147989 | Human | TME | 2 | 7389 | 10x Genomics | Primary | 32601337 |
| BCC_GSE123813_aPD1 | Human | Immunotherapy | 11 | 52884 | 10x Genomics | Metastatic | 31359002 |
| BLCA_GSE130001 | Human | TME | 2 | 4129 | 10x Genomics | Primary | 32111252 |
| BLCA_GSE145281_aPD1 | Human | Immunotherapy | 14 | 58683 | Smart-seq2 | Metastatic | 32405063 |
| BRCA_GSE110686 | Human | TME | 2 | 6035 | 10x Genomics | Primary, Metastatic | 29942092 |
| BRCA_GSE114727_10X | Human | TME | 3 | 28678 | 10x Genomics | Primary | 29961579 |
| BRCA_GSE114727_inDrop | Human | TME | 8 | 19676 | inDrop | Primary | 29961579 |
| BRCA_GSE136206_mouse_aPD1aCTLA4 | Mouse | Immunotherapy | NA | 27532 | 10x Genomics | Primary | 31730857 |
| BRCA_GSE138536 | Human | TME | 8 | 1902 | Smart-seq2 | Primary | 31974247 |
| BRCA_GSE143423 | Human | TME | 2 | 4375 | 10x Genomics | Metastatic | bioRxiv |
| BRCA_SRP114962 | Human | TME | 8 | 2472 | SNRS | Primary | 29681456 |
| CHOL_GSE125449_aPD1aPDL1aCTLA4 | Human | Immunotherapy | 10 | 5761 | 10x Genomics | Primary | 31588021 |
| CLL_GSE111014 | Human | TME | 4 | 30106 | 10x Genomics | Primary | 31996669 |
| CLL_GSE125881 | Human | Immunotherapy | 4 | 60528 | 10x Genomics | Primary | 31924795 |
| COAD_GSE108989 | Human | TME | 12 | 11125 | Smart-seq2 | Primary | 30479382 |
| COAD_GSE112865_mouse_aPD1 | Mouse | Immunotherapy | NA | 4454 | 10x Genomics | Primary | 30613266 |

| | | | | | | | |
|---|---|---|---|---|---|---|---|
| COAD_GSE120909_mouse_aPD1 | Mouse | Immunotherapy | NA | 1881 | Smart-seq | Primary | 30389797 |
| COAD_GSE122969_mouse_aPD1aTIM3 | Mouse | Immunotherapy | NA | 5457 | 10x Genomics | Primary | 30635236 |
| COAD_GSE136394 | Human | Immunotherapy | 5 | 67171 | 10x Genomics | Primary, Metastatic | 31484655 |
| COAD_GSE139555 | Human | TME | 2 | 10112 | 10x Genomics | Primary | 32103181 |
| COAD_GSE146771_10X | Human | TME | 10 | 43817 | 10x Genomics | Primary | 32302573 |
| COAD_GSE146771_Smartseq2 | Human | TME | 10 | 10468 | Smart-seq2 | Primary | 32302573 |
| GBM_GSE102130 | Human | TME | 6 | 3321 | Smart-seq2 | Primary | 29674595 |
| GBM_GSE103224 | Human | TME | 8 | 17185 | Microwell | Primary | 30041684 |
| GBM_GSE131928_10X | Human | TME | 9 | 13553 | 10x Genomics | Primary | 31327527 |
| GBM_GSE131928_Smartseq2 | Human | TME | 28 | 7930 | Smart-seq2 | Primary | 31327527 |
| GBM_GSE135437 | Human | TME | 19 | 12559 | mCEL-Seq2 | Primary, Metastatic | 31740814 |
| GBM_GSE138794 | Human | TME | 9 | 18458 | 10x Genomics | Primary | 31554641 |
| GBM_GSE139448 | Human | TME | 3 | 12152 | 10x Genomics | Primary | 32004492 |
| GBM_GSE141982 | Human | TME | 2 | 5263 | 10x Genomics | Primary | 32105316 |
| GBM_GSE148842 | Human | TME | 7 | 111397 | Microwell | Primary | bioRxiv |
| GBM_GSE70630 | Human | TME | 6 | 4347 | Smart-seq2 | Primary | 27806376 |
| GBM_GSE84465 | Human | TME | 4 | 3533 | Smart-seq2 | Primary | 29091775 |
| GBM_GSE89567 | Human | TME | 10 | 6341 | Smart-seq2 | Primary | 28360267 |
| HNSC_GSE103322 | Human | TME | 18 | 5902 | Smart-seq2 | Primary | 29198524 |
| HNSC_GSE139324 | Human | TME | 26 | 130721 | 10x Genomics | Primary | 31924475 |
| KIRC_GSE111360 | Human | TME | 2 | 23130 | 10x Genomics | Primary | 30550791 |
| KIRC_GSE145281_aPD1 | Human | Immunotherapy | 4 | 44220 | 10x Genomics | Metastatic | 32103181 |
| KIRC_GSE139555 | Human | TME | 3 | 49907 | 10x Genomics | Primary | 32405063 |

| | | | | | | | |
|---|---|---|---|---|---|---|---|
| LIHC_GSE125449_aPDL1aCTLA4 | Human | Immunotherapy | 9 | 3834 | 10x Genomics | Primary | 31588021 |
| LIHC_GSE140228_10X | Human | TME | 5 | 62530 | 10x Genomics | Primary | 31675496 |
| LIHC_GSE140228_Smartseq2 | Human | TME | 6 | 7074 | Smart-seq2 | Primary | 31675496 |
| LIHC_GSE98638 | Human | TME | 6 | 5059 | Smart-seq2 | Primary | 28622514 |
| MB_GSE119926 | Human | TME | 25 | 7745 | Smart-seq2 | Primary, Metastatic | 31341285 |
| MCC_GSE117988_aPD1aCTLA4 | Human | Immunotherapy | 1 | 10134 | 10x Genomics | Metastatic | 30250229 |
| MCC_GSE118056_aPDL1 | Human | Immunotherapy | 1 | 11024 | 10x Genomics | Primary | 30250229 |
| MM_GSE117156 | Human | TME | 14 | 24918 | MARS-seq | Primary | 30523328 |
| MM_GSE141299 | Human | TME | 7 | 16840 | 10x Genomics | Primary | NA |
| NET_GSE140312 | Human | TME | 1 | 3158 | 10x Genomics | Primary, Metastatic | 32054662 |
| NHL_GSE128531 | Human | TME | 9 | 30497 | 10x Genomics | Primary | 31010835 |
| NSCLC_EMTAB6149 | Human | TME | 8 | 40218 | 10x Genomics | Primary | 29988129 |
| NSCLC_GSE117570 | Human | TME | 4 | 11453 | 10x Genomics | Primary | 31033233 |
| NSCLC_GSE127465 | Human | TME | 7 | 31179 | Smart-seq2 | Primary | 30979687 |
| NSCLC_GSE127471 | Human | TME | 1 | 1108 | 10x Genomics | Primary | 31061481 |
| NSCLC_GSE131907 | Human | TME | 44 | 203298 | 10x Genomics | Primary, Metastatic | 32385277 |
| NSCLC_GSE139555 | Human | TME | 6 | 78829 | 10x Genomics | Primary | 32103181 |
| NSCLC_GSE143423 | Human | TME | 3 | 12193 | 10x Genomics | Metastatic | bioRxiv |
| NSCLC_GSE99254 | Human | TME | 14 | 12346 | Smart-seq2 | Primary | 29942094 |
| OV_GSE115007 | Human | TME | 1 | 6000 | 10x Genomics | Primary | 29967419 |
| OV_GSE118828 | Human | TME | 9 | 1909 | Smart-seq2 | Primary, Metastatic | 30383866 |
| PAAD_CRA001160 | Human | TME | 35 | 57443 | 10x Genomics | Primary | 31273297 |
| PAAD_GSE111672 | Human | TME | 3 | 6122 | inDrop | Primary | 31932730 |

| | | | | | | | |
|---|---|---|---|---|---|---|---|
| PBMC_30K_10X | Human | Normal | 1 | 29079 | 10x Genomics | Normal | 10X Genomics |
| PBMC_60K_10X | Human | Normal | 1 | 63628 | 10x Genomics | Normal | 10X Genomics |
| PBMC_8K_10X | Human | Normal | 1 | 8488 | 10x Genomics | Normal | 10X Genomics |
| SARC_GSE119352_mouse_aPD1aCTLA4 | Mouse | Immunotherapy | NA | 13789 | 10x Genomics | Primary | 30343900 |
| SCC_GSE123813_aPD1 | Human | Immunotherapy | 4 | 25891 | 10x Genomics | Metastatic | 31359002 |
| SKCM_GSE115978_aPD1 | Human | Immunotherapy | 31 | 7186 | Smart-seq2 | Primary, Metastatic | 30388455 |
| SKCM_GSE120575_aPD1aCTLA4 | Human | Immunotherapy | 48 | 16291 | Smart-seq2 | Metastatic | 30388456 |
| SKCM_GSE123139 | Human | TME | 25 | 35494 | MARS-seq | Primary, Metastatic | 30595452 |
| SKCM_GSE139249 | Human | TME | 4 | 39884 | 10x Genomics | Metastatic | 31801909 |
| SKCM_GSE148190 | Human | TME | 3 | 27834 | 10x Genomics | Metastatic | 32539073 |
| SKCM_GSE72056 | Human | TME | 19 | 4645 | Smart-seq2 | Metastatic | 27124452 |
| STAD_GSE134520 | Human | TME | 13 | 41554 | 10x Genomics | Primary | 31067475 |
| UCEC_GSE139555 | Human | TME | 3 | 12758 | 10x Genomics | Primary | 32103181 |
| UVM_GSE139829 | Human | TME | 11 | 103703 | 10x Genomics | Primary, Metastatic | 31980621 |

**Supplementary Table S2**

The criteria of cell clustering for setting the number of principal components and resolution based on the number of cells.

| Number of cellls | Number of PCs | Resolution |
|---|---|---|
| 1000 - 5000 | 15 | 0.6 |
| 5001 - 10000 | 20 | 1 |
| 10001 - 40000 | 30 | 1 |
| 40001 - 80000 | 40 | 1 |
| 80001 - 150000 | 50 | 1 |
| 150001 - | 75 | 1 |

**Supplementary Table S3**
Marker genes collected and curated for major-lineage level of cell-type annotation.

| Major-lineage cell type | Markers |
|---|---|
| B | ABCB4, ADAM28, BACH2, BANK1, BCL7A, BEND5, BLK, BRAF, CD180, CD19, CD1C, CD22, CD37, CD69, CD72, CD79A, CD79B, CR2, CXCR5, EAF2, FAIM3, FCER2, FCGR2B, FCRL2, FRK, GPR18, GUSBP11, HHEX, HLA-DOB, IGHD, IGHM, IGKC, IGLL3P, IL4R, IRF8, KIAA0226L, LINC00921, LTB, LY86, MEP1A, MICAL3, MS4A1, NIPSNAP3B, NMBR, P2RX5, P2RY14, PNOC, PSG2, PTPRCAP, RALGPS2, RASGRP2, SELL, SIK1, SLC12A1, SPIB, STAP1, TCL1A, UGT1A8, VPREB3, ZNF286A, AIM2, ALOX5, CCR6, CD27, CLCA3P, DENND5B, FAM65B, GNG7, IFNA10, IL7, MBL2, NPIPB15, SIT1, SP140, TMEM156, TNFRSF13B, TNFRSF17, TRAF4, ZBTB32 |
| CD4Tconv | ACAP1, ANKRD55, ATHL1, BCL11B, CCR7, CD2, CD247, CD27, CD3D, CD3G, CD40LG, CD7, CXorf57, DPP4, DSC1, EPHA1, FAIM3, FLJ13197, FLT3LG, GAL3ST4, GALR1, GPR1, GRAP2, GZMM, ICOS, IL7R, ITK, LAT, LCK, LEF1, LIME1, LTB, LY9, MAP4K1, MAP4K2, MAP9, RASGRP2, RPL3P7, SERGEF, SH2D1A, SIRPG, TCF7, TRAC, TRAT1, TRAV13-1, TRBC1, UBASH3A, VILL, WNT7A, ZAP70, ZNF204P, ZNF324, CCL5, CCR6, CD28, CD3E, CD4, CD6, CD69, CD96, CTLA4, CTSW, DGKA, EPB41, ETS1, FBXL8, GPR171, GPR25, GZMA, GZMK, KLRB1, NKG7, PBXIP1, PTGER2, PTPRCAP, RASA3, RCAN3, RPL10L, ST8SIA1, TRAV13-2, TRAV21, TRAV8-6, TRAV9-2, ZFP36L2, CCL20, CDC25A, CSF2, CXCL13, GPR19, GZMB, IFNG, IL12RB2, IL17A, IL26, IL2RA, IL3, IL4, IL9, LAG3, LTA, ORC1, PMCH, RRP9, SKA1, TNFRSF4, TNIP3, CHI3L2, CXCR5, FOSB, FZD3, ICA1, IL21, PASK, PDCD1, PVRIG, RGS1, SIK1, SLC7A10, TRIB2, TSHR, ZBTB10 |
| CD8T | BCL11B, CCL5, CD2, CD247, CD27, CD3D, CD3E, CD3G, CD6, CD69, CD7, CD8A, CD8B, CD96, CRTAM, CST7, CTSW, DPP4, DSC1, DUSP2, FAIM3, FLT3LG, GNLY, GPR171, GRAP2, GZMA, GZMB, GZMH, GZMK, GZMM, ICOS, IGKC, IL7R, ITK, KLRB1, KLRC3, KLRC4, KLRD1, KLRF1, KLRK1, LAG3, LCK, LEF1, LIME1, LTB, LY9, MAP4K1, MAP9, NCR3, NKG7, PIK3IP1, PRF1, PTGDR, PTPRCAP, PVRIG, RASA3, RPL3P7, SH2D1A, SIRPG, TCF7, TRAC, TRAT1, TRAV12-2, TRAV13-1, TRBC1, TRDC, UBASH3A, ZAP70 |
| CD8Tex | BCL11B, CCL5, CD2, CD247, CD27, CD3D, CD3E, CD3G, CD6, CD69, CD7, CD8A, CD8B, CD96, CRTAM, CST7, CTSW, DPP4, DSC1, DUSP2, FAIM3, FLT3LG, GNLY, GPR171, GRAP2, GZMA, GZMB, GZMH, GZMK, GZMM, ICOS, IGKC, IL7R, ITK, KLRB1, KLRC3, KLRC4, KLRD1, KLRF1, KLRK1, LAG3, LCK, LEF1, LIME1, LTB, LY9, MAP4K1, MAP9, NCR3, NKG7, PIK3IP1, PRF1, PTGDR, PTPRCAP, PVRIG, RASA3, RPL3P7, SH2D1A, SIRPG, TCF7, TRAC, TRAT1, TRAV12-2, TRAV13-1, TRBC1, TRDC, UBASH3A, ZAP70, PDCD1, CTLA4, TIGIT, HAVCR2 |
| DC | AIF1, ALOX15, C1orf54, CCDC102B, CCL13, CCL17, CCL18, CCL22, CD1A, CD1B, CD1C, CD1E, CD209, CD33, CD68, CLEC10A, CLEC4A, CLEC7A, CLIC2, DHRS11, EGR2, FAM198B, FCER1A, FCER2, FLVCR2, FPR3, FZD2, HLA-DQA1, IGSF6, MMP12, NCF2, PLA2G7, PPFIBP1, RNASE6, SCN9A, SLAMF8, SLC15A3, TMEM255A, TREM2, ARHGAP22, BIRC3, CCL1, CCL19, CCL20, CCL5, CCL8, CCR7, CD80, CD86, CHST7, CXCL10, CXCL11, CYP27A1, DHX58, EBI3, ETV3, HESX1, HTR2B, IDO1, IFI44L, IL12B, IL2RA, KYNU, LAMP3, MAP3K13, MSC, NR4A3, PDCD1LG2, PLA1A, PTGIR, RASSF4, RSAD2, SIGLEC1, SLC2A6, SLCO5A1, ST3GAL6, TNFAIP6, TNFRSF11A, TNFRSF4 |
| Endothelial | PECAM1, VWF, ENG |

| | |
|---|---|
| Fibroblasts | FAP, PDPN, MMP2, PDGFRA, THY1, MMP11, PDGFRL, TGFB3, COL1A2, DCN, COL3A1, COL6A1 |
| Mast | ATP8B4, BMP2K, BPI, C3AR1, CD33, CEACAM8, CLC, CMA1, CPA3, CRISP3, CTSG, FAM124B, FAM174B, FCER1A, GFI1, HDC, HPGDS, IL18R1, LTC4S, MS4A2, MS4A3, MYB, NOX3, NTRK1, P2RX1, P2RY14, PAQR5, PRG2, RAB27B, RGS13, SEPT8, SLC12A8, ST8SIA1, STAP1, STXBP6, TPSAB1, CCL1, CCL20, CCL4, CSF2, CXCL3, GZMB, HOXA1, IL1A, IL1B, IL1RL1, IL3, IL5, LINC00597, MARCH3, TEC |
| Mono/Macro | AIF1, APOBEC3A, AQP9, ASGR1, ASGR2, BST1, C5AR1, CCR2, CD1D, CD33, CD68, CDA, CFP, CHST15, CLEC4A, CLEC7A, CREB5, CSF3R, FAM198B, FCN1, FES, FOSB, FPR1, FZD2, HCK, HK3, HNMT, HPSE, IGSF6, LILRA2, LILRA3, LILRB2, LST1, MEFV, MNDA, MS4A6A, NCF2, NFE2, NLRP3, NOD2, P2RY13, PADI4, RNASE2, RNASE6, S100A12, SLC15A3, TLR2, TLR7, TLR8, UPK3A, VNN1, VNN2, ACP5, ADAMDEC1, BHLHE41, CCDC102B, CCL18, CCL22, CCL7, CHI3L1, COL8A2, CSF1, CXCL3, CXCL5, CYP27A1, DCSTAMP, GPC4, MARCO, MMP9, PLA2G7, PPBP, QPCT, SLAMF8, SLC12A8, TNFSF14, TREM2, ACHE, APOL3, APOL6, ARRB1, CCL19, CCL5, CCL8, CCR7, CD38, CD40, CLIC2, CXCL10, CXCL11, CXCL13, CXCL9, CYP27B1, DHX58, EBI3, GGT5, HESX1, IDO1, IFI44L, IL2RA, KIAA0754, KYNU, LAG3, LAMP3, PLA1A, PTGIR, RASSF4, RSAD2, SIGLEC1, SLAMF1, SLC2A6, SOCS1, TNFAIP6, TNIP3, TRPM4, ALOX15, CCL13, CCL14, CCL23, CD209, CD4, CLEC10A, CRYBB1, FRMD4A, GSTT1, HRH1, HTR2B, NME8, NPL, PDCD1LG2, RENBP, WNT5B |
| Myofibroblasts | ACTA2, MCAM, MYLK, MYL9, IL6, PDGFA |
| Neutrophils | AIF1, APOBEC3A, AQP9, BTNL8, C5AR1, CAMP, CASP5, CCR3, CDA, CEACAM3, CFP, CHI3L1, CHST15, CLC, CREB5, CSF3R, CXCR1, CXCR2, DPEP2, EMR2, EMR3, FAM212B, FCGR3B, FFAR2, FPR1, FPR2, GPR97, HAL, HSPA6, IGSF6, IL18RAP, LILRA2, LILRB2, LST1, MAK, MEFV, MGAM, MMP25, MNDA, MXD1, NCF2, NFE2, P2RY13, P2RY14, PADI4, PGLYRP1, PLEKHG3, QPCT, REPS2, S100A12, STEAP4, TLR2, TLR8, TNFAIP6, TNFRSF10C, TREM1, TREML2, VNN1, VNN2, VNN3 |
| NK | BPI, CAMP, CCL5, CD160, CD2, CD244, CD247, CD7, CD96, CDHR1, CEACAM8, CST7, CTSW, DEFA4, ELANE, GFI1, GNLY, GZMA, GZMB, GZMH, GZMK, GZMM, IL12RB2, IL18R1, IL18RAP, IL2RB, KIR2DL1, KIR3DL2, KLRB1, KLRC3, KLRC4, KLRD1, KLRF1, KLRK1, LCK, MGAM, MS4A3, NAALADL1, NKG7, NME8, PLEKHF1, PRF1, PRR5L, PTGDR, PTPRCAP, PVRIG, S1PR5, SH2D1A, TBX21, TEP1, TRBC1, TRDC, TTC38, TXK, ZAP70, ZNF135, APOBEC3G, APOL6, CCL4, CCND2, CD69, CDK6, CSF2, DPP4, FASLG, GPR171, GPR18, GRAP2, IFNG, KIR2DL4, KIR2DS4, LTA, LTB, NCR3, OSM, PTGER2, SOCS1, TNFSF14 |
| pDC | AIF1, ALOX15, C1orf54, CCpDC102B, CCL13, CCL17, CCL18, CD1A, CD1B, CD1C, CD1E, CD33, CD68, CLEC10A, CLEC4A, CLEC7A, CLIC2, DHRS11, EGR2, FAM198B, FCER1A, FCER2, FLVCR2, FPR3, FZD2, HLA-DQA1, IGSF6, MMP12, NCF2, PLA2G7, PPFIBP1, RNASE6, SCN9A, SLAMF8, SLC15A3, TMEM255A, TREM2, CLEC4C, IRF7, LILRB4, Siglech, ARHGAP22, BIRC3, CCL1, CCL19, CCL20, CCL5, CCL8, CCR7, CD80, CD86, CHST7, CXCL10, CXCL11, CYP27A1, DHX58, EBI3, ETV3, HESX1, HTR2B, IDO1, IFI44L, IL12B, IL2RA, KYNU, LAMP3, MAP3K13, MSC, NR4A3, PpDCD1LG2, PLA1A, PTGIR, RASSF4, RSAD2, SIGLEC1, SLC2A6, SLCO5A1, ST3GAL6, TNFAIP6, TNFRSF11A, TNFRSF4 |
| Plasma | ABCB9, AMPD1, ANGPT4, ATXN8OS, C11orf80, CCR10, CD27, CD38, CD79A, DENND5B, EAF2, FCRL2, GNG7, GPR25, GUSBP11, HIST1H2AE, HIST1H2BG, HLA-DOB, IGHD, IGHE, IGHM, IGKC, IGLL3P, KCNA3, KCNG2, LIME1, LOC100130100, MAN1A1, MANEA, MAST1, MROH7, MZB1, P2RX5, PAX7, PDK1, PNOC, RASGRP3, REN, RGS13, RPL3P7, SIK1, SPAG4, ST6GALNAC4, TGM5, TMEM156, TNFRSF17, UGT2B17, ZBP1, ZNF165 |

| | |
|---|---|
| TMKI67 | BCL11B, CD2, CD247, CD27, CD28, CD3D, CD3E, CD3G, CD6, CD7, CD8A, CD8B, CD96, CXCR6, FLT3LG, FYN, GIMAP4, GPR171, GZMK, GZMM, ICOS, ITK, LCK, LIME1, PRKCH, PSTPIP1, SH2D1A, SIRPG, TNFRSF9, TRAC, TRAT1, TRBC1, TRBC2, UBASH3A, ZAP70, AURKA, BIRC5, BUB1, CCNA2, CCNB1, CDC20, CDK1, CDKN3, FEN1, HMGB2, MCM2, MCM5, MCM6, MYBL2, NUSAP1, PCNA, PLK1, TOP2A, ZWINT |
| Treg | BARX2, BCL11B, CD2, CD247, CD27, CD28, CD3D, CD3E, CD3G, CD4, CD5, CD6, CD70, CD96, CEMP1, CLEC2D, CTLA4, DGKA, DPP4, EFNA5, FOXP3, FRMD8, GPR1, GPR171, GPR19, GZMM, HIC1, HMGB3P30, ICOS, IL2RA, IL2RB, ITK, KIRREL, LAIR2, LCK, LILRA4, LOC126987, LTB, MAP4K1, MBL2, NPAS1, NTN3, PCDHA5, PLCH2, PMCH, PTGIR, PTPRG, RCAN3, RYR1, SEC31B, SEPT5, SH2D1A, SIRPG, SIT1, SKAP1, SPOCK2, SSX1, TRAC, TRAT1, TRAV9-2, TRBC1, TYR, UBASH3A, ZAP70 |

**Supplementary Table S4**
Marker genes collected and curated for minor-lineage level of cell-type annotation.

| Minor-lineage cell type | Markers |
|---|---|
| CD4Teff | CD3D, CD3E, CD3G, CD4, CTSW, CX3CR1, GNLY, GZMH, KLRG1, NKG7, PRF1, S1PR1, S1PR5, TBX21 |
| CD4Tn | CCR7, CD27, CD28, CD3D, CD3E, CD3G, CD4, LEF1, RUNX3, S1PR1, SELL, TCF7, ZBTB7B |
| CD8Tcm | BCL6, BCL6B, BMI1, CCL5, CCR7, CD27, CD28, CD3D, CD3E, CD3G, CD44, CD8A, CD8B, CXCR3, EOMES, GPR183, GZMA, GZMK, IL15RA, IL2RB, IL2RG, IL7R, MBD2, PRF1, S1PR1, SELL, TRD, TRG |
| CD8Teff | CD3D, CD3E, CD3G, CD8A, CD8B, CX3CR1, EOMES, FCGR3A, FGFBP2, FOS, GZMH, IFNG, JUN, KLRG1, PRF1, S1PR1, S1PR5, TBX21, TNF |
| CD8Tem | CCR7, CD3D, CD3E, CD3G, CD44, CD8A, CD8B, CXCR3, CXCR4, FGFBP2, GZMK, IL15RA, IL2RB, IL2RG, IL7R, KLRD1, KLRG1, PRDM1, TRD, TRG |
| CD8Tn | CCR7, CD27, CD28, CD3D, CD3E, CD3G, CD8A, CD8B, IL7R, LEF1, RUNX3, S1PR1, SELL, TCF7, ZBTB7B |
| cDC1 | BATF3, BTLA, CADM1, CD226, CLEC9A, DPP4, ID2, IRF8, THBD, TLR3, XCR1, CCL22, FLT3, CD8A, CD207, ITGAE |
| cDC2 | CCL22, CD209, CCL17, HSD11B1, CCL13, PPFIBP2, NPR1, CD1B, VASH1, F13A1, CD1E, MMP12, FABP4, CLEC10A, SYT17, MS4A6A, CTNS, GUCA1A, CARD9, ABCG2, CD1A, PPARG, RAP1GAP, SLC7A8, GSTT1, PDXK, FZD2, CSF1R, HS3ST2, CH25H, LMAN2L, SLC26A6, BLVRB, NUDT9, PREP, TM7SF4, TACSTD2, CD1C, CCL1, EBI3, INDO, LAMP3, OAS3, IL3RA |
| M1 | SIGLEC1, CCR5, CD33, CD40, CD63, CD68, CD80, CD86, CLEC7A, CXCL10, CXCL11, CXCL2, CXCL9, ENG, FCGR1A, FCGR1B, FCGR1C, FCGR2A, FCGR2B, FCGR3A, FCGR3B, FUT4, IDO, LILRA4, MerTK, NOS2, SOCS1, SOCS3, TLR2, TLR4, TREM1, FCER1G, FCER2A, HLA-DRA |
| M2 | SIGLEC1, ARG1, CCL22, CCR5, CD163, CD200R1, CD209, CD33, CD40, CD63, CD68, CD80, CD86, CXCL10, CXCL11, CXCL2, CXCL9, ENG, FCER2, FUT4, ITGAM, JUN, LILRA4, MRC1, MSR1, PPARG, TGM2, TLR2, TLR4, TREM1, TREM2, FCER1G, FCER2A, HLA-DRA |
| MAIT | CD3D, CD3E, CD3G, CD8A, CD8B, KLRB1, NCR3, RORA, RORC, SLC4A10, ZBTB16 |
| Monocyte | SIGLEC1, CCR5, CD14, CD33, CD40, CD63, CD68, CD80, CD86, CD93, CSF1R, CXCL10, CXCL11, CXCL2, CXCL9, ENG, FCGR3A, FUT4, ITGA5, ITGAM, ITGAX, LILRA4, S100A9, TLR2, TLR4, TREM1, S100A8 |

| | |
|---|---|
| pDC | CD123, CD14, CD19, CD1A, CLEC4C, DR6, FECR1, GZMB, HLA-DRA, IFNA1, IFNB1, IL10, IL3RA, IL6, ILT3, ILT7, IRF4, IRF7, IRF8, ITGAX, LILRB4, MS4A1, NCAM1, NRP1, PTPRC, Siglech, SPIB, TCF4, TLR7, TLR9, TNF, ZEB2, FLT3 |
| Plasma | ABCB9, AMPD1, ANGPT4, ATXN8OS, C11orf80, CCR10, CD27, CD38, CD79A, DENND5B, EAF2, FCRL2, GNG7, GPR25, GUSBP11, HIST1H2AE, HIST1H2BG, HLA-DOB, IGHD, IGHE, IGHM, IGKC, IGLL3P, KCNA3, KCNG2, LIME1, LOC100130100, MAN1A1, MANEA, MAST1, MROH7, MZB1, P2RX5, PAX7, PDK1, PNOC, RASGRP3, REN, RGS13, RPL3P7, SIK1, SPAG4, ST6GALNAC4, TGM5, TMEM156, TNFRSF17, UGT2B17, ZBP1, ZNF165 |
| Tfh | BCL6, BLTA, CD200, CD3D, CD3E, CD3G, CD4, CD40LG, CXCR3, CXCR5, ICA1, ICOS, IL21R, IL6ST, MAGEH1, PDCD1, PTPN13, SLAM, STAT3, TNFSF4, TOX, TOX2 |
| Th1 | BHLHE40, CCR1, CCR5, CD3D, CD3E, CD3G, CD4, CD94, CXCL13, CXCR3, CXCR6, GZMB, HAVCR2, ICOS, IFNG, IFNGR1, IFNGR2, IGFLR1, IL12RB1, IL12RB2, IL18R1, ITGAE, PDCD1, STAT1, STAT4, TBX21, TNFSF11 |
| Th17 | CAPG, CCR6, CD3D, CD3E, CD3G, CD4, CTSH, FURIN, IL23R, IL17A, IL1R1, IL1R2, IL26, ITGAE, KLRB1, RORA, RORC, STAT3 |
| Th2 | BHLHE41, CCR4, CCR8, CD3D, CD3E, CD3G, CD4, CXCR4, GATA3, GPR44, HAVCR1, IL17RB, IL1RL1, IL4R, MAF, PTGDR2, STAT6 |
| Treg | CD3D, CD3E, CD3G, CD4, CDC25B, CTLA4, FOXO1, FOXO3, FOXP3, IKZF2, IL10RA, IL2RA, RTKN2, S1PR4, STAT5A, STAT5B |
